# Diversification or collapse of self-incompatibility haplotypes as outcomes of evolutionary rescue

**DOI:** 10.1101/641613

**Authors:** Alexander Harkness, Emma E. Goldberg, Yaniv Brandvain

## Abstract

Self-incompatibility systems in angiosperms are exemplars of extreme allelic polymorphism maintained by long-term balancing selection. Pollen that shares an allele with the pollen recipient at the self-incompatibility locus is rejected, and this rejection favors rare alleles as well as preventing self-fertilization. Advances in molecular genetics reveal that an ancient, deeply conserved, and well-studied incompatibility system functions through multiple tightly linked genes encoding separate pollen-expressed F-box proteins and pistil-expressed ribonucleases. We show that certain recombinant haplotypes at the incompatibility locus can drive collapse in the number of incompatibility types. We use a modified evolutionary rescue model to calculate the relative probabilities of increase and collapse in number of incompatibility types given the initial collection of incompatibility haplotypes and the population rate of gene conversion. We find that expansion in haplotype number is possible when population size or the rate of gene conversion is large, but large contractions are likely otherwise. By iterating a Markov chain model derived from these expansion and collapse probabilities, we find that a stable haplotype number distribution in the realistic range of 10–40 is possible under plausible parameters. However, small or moderate-sized populations should be susceptible to substantial additional loss of haplotypes beyond those lost by chance during bottlenecks. The same processes that can generate many incompatibility haplotypes in large populations may therefore be crushing haplotype diversity in smaller populations.

## Introduction

Self-incompatibility (SI) is a common strategy by which plants ensure outcrossing, and it provides a classic example of extreme allelic polymorphism maintained by long-term balancing selection. An SI plant rejects self pollen, which is identified by a specificity phenotype encoded by a highly polymorphic self-incompatibility locus (S-locus). This type of inbreeding avoidance mechanism is widespread in plants: distinct non-homologous SI systems have been discovered in the Poaceae (Li et al. 1997), Papaveraceae (Foote et al. 1994), Solanaceae (McClure et al. 1989), Brassicaceae (Stein et al. 1991), and Asteraceae (Hiscock et al. 2003), and some form of SI is thought to be present in nearly 40% of plant species across 100 families (Igić et al. 2008). Rejection occurs when the pollen specificity matches the pistil specificity, which is likewise encoded by the S-locus. Pollen with a rare specificity has an advantage because it is less likely to encounter a pistil with a matching specificity and is thus less likely to be rejected. This advantage of rarity results in balancing selection, which maintains polymorphism at the S-locus by protecting S-locus alleles (S-alleles) from loss through drift (Wright 1939). While it is “fairly obvious that selection would tend to increase the frequency of any additional alleles that may appear” (Wright 1939), it is not obvious how novel S-alleles do appear. Molecular genetic understanding of SI has advanced rapidly, and modern theory to explain the diversification of S-alleles must account for what is now known about the structure of the S-locus. We develop a population genetic model of the expansion, collapse, and long-term evolution of S-allele number under the widespread solanaceous SI system.

The SI system originally discovered in *Nicotiana* (East and Mangelsdorf 1925) is particularly well-studied. Counts of 10–28 alleles have been directly observed in several other species in Solanaceae, and *Physa¡is crassifolia* has been estimated to harbor as many as 44 (Lawrence 2000). In the solanaceous SI system, each haploid pollen grain carries one allele at the S-locus, and the pollen is rejected if its allele matches either of those carried by the diploid pollen recipient (East and Mangelsdorf 1925). Similar systems in the Plantaginaceae and the distantly related Rosaceae have been found to be homologous, which implies that the solanaceous system was present in the common ancestor of the asterids and rosids (Igić and Kohn 2001; Steinbachs and Holsinger 2002), estimated to have lived about 120 Mya (Tank et al. 2015). S-alleles appear to be very long-lived, consistent with long-term balancing selection. A given allele is often more closely related to an allele in another species or even another genus than it is to other alleles in the same population, which is consistent with the alleles having persisted since the common ancestor of those genera (Igic and Kohn 2001). Theory for the origin and fate of S-alleles can help us understand the forces shaping allelic diversity within and across species.

### Conceptual challenges

A series of influential models investigated how a system of many incompatibility alleles could arise, but only very recently have theorists incorporated assumptions fully consistent with the contemporary understanding of the molecular genetics of self-incompatibility. Initially, the answer seemed simple. Wright (1939) showed that each fully functional S-allele is under negative frequency-dependent selection because pollen carrying rare alleles is compatible with a larger fraction of the population. This classic model predicts the number of S-alleles maintained in a population as a balance between their gain by mutation or immigration and their loss by drift. Charlesworth and Charlesworth (1979) showed that functional S-alleles can invade an initially SI population when inbreeding depression is high and that, conversely, S-locus mutations abolishing SI can invade when inbreeding depression is low. This model provided a plausible hypothesis for the origin of SI. However, these one-gene models of incompatibility were inconsistent with the separate pistil-(McClure et al. 1989) and pollen-expressed (Lai et al. 2002; Entani et al. 2003; Sijacic et al. 2004) products that were later discovered. Uyenoyama et al. (2001) and Gervais et al. (2011) showed that if each specificity is instead controlled by two tightly linked pollen- and pistil-function loci, new S-alleles can be generated through a self-compatible intermediate, but one or more S-alleles are also lost for many parameter combinations. Sakai (2016) additionally considered that new alleles arise from old alleles and showed that if each pollen-function mutation is more likely to be rejected by the pistil-function allele that was on its initial genetic background, then the number of S-alleles can increase from zero into the range observed in nature. A critical common element among all these models is the assumption that SI functions through self-recognition. In self-recognition systems, pollen is accepted by default, but the pistil recognizes and rejects self pollen based on a match between complementary pistil- and pollen-products.

However, the solanaceous SI system is now known to function not through self-recognition, but rather through a form of nonself-recognition (Kubo et al. 2010). In nonself-recognition systems, pollen is rejected by default, but complementary pollen- and pistil-products react to prevent rejection of cross pollen. Results from models of self-recognition systems may not be directly applicable to nonself-recognition. Despite the ubiquity of solanaceous SI, few models of S-allele evolution explicitly incorporate its nonself-recognition system. Fujii et al. (2016) proposed that a pollen-function mutation complementary to a novel pistil-function mutation could arise on a single background and then spread to other backgrounds through gene conversion. Bod’ová et al. (2018) enumerated several possible evolutionary pathways to new S-alleles, and their results also showed that the pathway hypothesized by Fujii et al. (2016) could either expand or reduce S-allele number. An explanation of the long-term trajectory of S-allele number must therefore account not only for the steps needed to create new fully-functioning alleles, but also for dynamics that can lead to their loss.

A stochastic process involving the risk of collapse raises radically different questions than one of continual growth. Are S-allele numbers repeatedly expanding and contracting, or do they reach equilibria where changes are rare? We may sometimes observe changes in progress if they are frequent, but infrequent changes may be buried in the past. When a contraction occurs, is it usually small or catastrophic? Large contractions, followed by re-expansion, should result in more turnover in alleles and fewer shared alleles among species than would small contractions. Since contraction is possible whether SI is maintained throughout (Bod’ová et al. 2018) or is temporarily lost in self-compatible intermediates (Uyenoyama et al. 2001; Gervais et al. 2011; Bod’ová et al. 2018), neither pathway is unidirectional. Is the pathway maintaining SI still sufficient for expansion, or are self-compatible intermediates necessary? If new proto-S-alleles are segregating in natural populations, are they self-compatible, -incompatible, or both? Answers to these questions will aid interpretation of S-allele genealogies, which document both similarities and differences between species in their complement of S-alleles. For example, the large historical contraction in S-allele number in the common ancestor of *Physalis* and *Witheringia* has been interpreted as a demographic bottleneck (Paape et al. 2008), but a firmer theoretical investigation could assess whether a collapse restricted to the S-locus is a plausible alternative explanation.

We use an evolutionary rescue model to approximate the relative probabilities of collapse and expansion in S-allele number as well as the distribution of collapse magnitudes. We find a negative relationship between initial S-allele number and the probability of further expansion. By constructing and iterating a Markov chain out of these contraction and expansion probabilities, we find stable long-term distributions of haplotype number. Realistic S-allele numbers (20–40) are possible through this process alone for large but plausible values of population size and rate of gene conversion. However, contractions can be very large when they occur, often eliminating the majority of S-alleles. These results suggest that contractions in S-allele number are ubiquitous unless new S-alleles are prevented from invading in the first place, as might occur if they cause pollen limitation.

Our results point to three major predictions. First, we predict that collapse is very likely when many existing S-alleles are pre-adapted to being compatible with novel S-alleles. This is because the pre-adapted alleles eliminate their competitors when a novel specificity arises, thereby reducing the total number of surviving alleles. Second, we predict that intraspecific variation in S-allele number can sometimes be explained by an invasion of a runaway self-incompatible allele rather than a bottleneck or a transition to self-compatibility. We should therefore expect to find allopatric populations with large disparities in S-allele number, and we should expect the population with the smaller number to harbor an S-allele that, if introduced into the other population, would trigger a collapse. Third, populations that have recently undergone such a collapse should be characterized by self-incompatibility but low S-allele diversity, and they should show a signature of a selective sweep at the S-locus but greater polymorphism outside it.

## Model and Results

The risk of contraction in S-allele number in the solanaceous incompatibility system is rooted in the system’s biological details. This system involves two kinds of complementary products: pistil-expressed ribonucleases (RNases) (McClure et al. 1989) and pollen-expressed F-box proteins (Lai et al. 2002; Entani et al. 2003; Sijacic et al. 2004). Together, these products achieve a form of collaborative nonself-recognition first described by Kubo et al. (2010). Under this system, each “allele” at the S-locus is actually a tightly linked haplotype containing one RNase gene and a collection of paralogous F-box genes. The RNase gene is highly polymorphic, and each functionally distinct haplotype possesses a different RNase allele. Nonself pollen is recognized through the complement of F-box proteins it expresses. Each F-box paralog produces a functionally distinct product, and each of these products is capable of detoxifying one or more forms of RNase. Pollen is only successful if it expresses both of the two F-box proteins to match the diploid pollen recipient’s two RNase alleles. Self-fertilization is prevented because each fully functional haplotype lacks a functional copy of the F-box gene that corresponds to the RNase on the same haplotype, and so each pollen grain necessarily lacks one of the two F-box proteins required to fertilize the plant that produced it. Rejection is not restricted to self-pollination: any two individuals that share one haplotype will reject half of each other’s pollen, while individuals that share both haplotypes are completely incompatible. Rejection is therefore more likely between closely related individuals. Tight genetic linkage across all components of the S-locus reduces the probability that recombination will cause a haplotype to lose a functional F-box paralog (reducing its siring opportunities) or gain the F-box paralog that detoxifies its own RNase (inducing self-compatibility).

This system presents two novel challenges that are absent in self-recognition systems. The first is a “chicken-egg problem.” A novel F-box specificity alone is at best neutral because it detoxifies an RNase that does not yet exist. A novel RNase alone is deleterious because it degrades all pollen and renders the plant ovule-sterile. Both of these mutations must invade in order to generate a new, fully functional S-haplotype. Second, for every novel RNase that arises, the corresponding F-box specificity must appear on every other haplotypic background in order to restore cross-compatibility among all haplotypes. Building on the hypothesis for expansion of haplotype number proposed by Fujii et al. (2016), Bod’ová et al. (2018) showed how cross-compatibility could be restored among all incompatibility classes after novel RNase and F-box mutations have invaded. If an initially neutral F-box mutation already exists when its complementary RNase mutation arises, the F-box mutation will then invade because it confers the advantage of compatibility with the new RNase. To restore full cross-compatibility among haplotypes, all haplotypes must acquire F-box paralogs complementary to all RNase alleles other than their own either through gene conversion (Fujii et al. 2016) or recurrent mutation (Bod’ová et al. 2018). This means that the haplotype bearing the RNase mutation must also acquire the F-box complementary to its ancestor. Once this occurs, the resulting haplotype is compatible with pollen recipients carrying all other haplotypes, but haplotypes still lacking the new F-box are not compatible with pollen recipients carrying the new RNase. As the RNase mutation increases in frequency, siring opportunities decrease for haplotypes that still lack the new F-box, and they are gradually driven extinct. If all doomed haplotypes acquire their missing F-box before they are lost, expansion has occurred. But if some doomed haplotypes go extinct, their RNase alleles are lost and contraction has occurred. Bod’ová et al. (2018) simulated the dynamics of this process along with several other expansion pathways and found that up to 14 haplotypes could be maintained.

Based on this biological background, we introduce a set of metaphors to make the interactions among haplotypes more intuitive. Each form of pistil-function RNase is a lock, and each form of pollen-function F-box protein is a key. A diploid plant codominantly expresses two different locks in its pistils. Each pollen grain expresses every paralogous key in its haploid genome, and these keys collectively form that pollen grain’s key ring. The pollen must unlock both of the pollen recipient’s locks in order to fertilize it, which requires keys to both locks. Each haplotype is self-incompatible because its key ring lacks the key to the lock on the same haplotype. Our model must explain two processes: the invasion of a novel lock and the evolution of the complementary key on other haplotypes.

### Initial conditions

We focus on a population without any self-compatible haplotypes. Several models of incompatibility evolution include a self-compatible intermediate (Uyenoyama et al. 2001; Gervais et al. 2011; Bod’ová et al. 2018), which can either lead to expansion or to total collapse of incompatibility. Self-compatible haplotypes do not lose siring opportunities as their frequency increases, so they are not under the same negative frequency-dependence as functional incompatibility haplotypes. If inbreeding depression is low, self-compatibility is often universally superior to self-incompatiblity, and the population will eventually become completely self-compatible at equilibrium (Uyenoyama et al. 2001; Gervais et al. 2011; Bod’ová et al. 2018). We do not retread this well-established result. Our population can instead be interpreted as one in which inbreeding depression is so great that any self-compatible haplotypes produced by mutation or gene conversion are effectively instantly removed by selection, although we do not explicitly model inbreeding depression.

Evolution of a new incompatibility class requires both a new lock and a new key. A lock mutation without a corresponding key would cause ovule sterility to the mutant carrying it and would thus be quickly eliminated, but it would not otherwise affect the population. In contrast, a key to a not-yet-extant lock would be neutral as long as there were no cost of expressing it. There should therefore be conditionally neutral functional diversity among duplicate keys. If a series of unsuccessful lock mutations culminates in one that corresponds to an extant key, the final mutation will appear to have been anticipated by that key. We therefore model the key appearing before its complementary lock with the knowledge that this pair of mutations may have been preceded by a series of pairs that occurred in the “wrong” order. The new lock allele then arises from its ancestral lock allele on one of the existing haplotypes. Biologically, this corresponds to a mutation changing the specificity of one of the existing RNase alleles such that its product is now recognized by a different F-box protein and no longer by the old one. Note that there are two other possible results of gene conversion besides adding a key to a key ring. First, gene conversion may replace a functional F-box paralog with a non-functional or deleted allele: i.e., it may remove a key. Such a conversion event should reduce the siring success of the haplotype and be unconditionally deleterious. Second, gene conversion may induce self-compatibility by uniting a key with its complementary lock. This conversion event should also be deleterious if SI is maintained by intense inbreeding depression. These convertants certainly occur, but we do not track them because they should be removed rapidly. Note that our gene conversion parameter, which includes only conversions that add a key without inducing self-compatibility, is therefore smaller than the raw rate of functional gene conversion, which also includes the production of short-lived deleterious convertants. Rather than speculate on the empirical proportions of these kinds of conversion events, we simply vary the rate of key additions that maintain SI.

A major barrier to expansion without a self-compatible intermediate is that, unless all other haplotypes possess the new key, the new lock mutation will necessarily reduce the amount of compatible pollen the mutant plant receives. If this reduced pollen supply reduces the mutant’s seed set, the mutation will be eliminated by selection because it provides no countervailing advantage. However, plants in large populations with efficient pollination may be so pollen-saturated that even a small compatible proportion is sufficient to achieve full seed set. We model such a population because it provides a biologically plausible (though probably not general) scenario in which a self-incompatible lock mutant can persist. Large populations are also the most conducive to expansion for reasons discussed below, so the absence of pollen limitation of seed set may be most realistic for the populations in which expansion is most likely. Strictly, though, the lock mutation can be neutral even if the mutant suffers some pollen limitation: it must merely suffer no more pollen limitation than average.

We consider a population with n initial complete haplotypes, each at frequency 1/*n* (Wright 1939). When one new lock mutation appears, we identify five classes of haplotypes (summarized in fig. 1). The mutation of the new lock is the first in a four-step evolutionary pathway to expansion or collapse (fig. 2). First, there are nL haplotypes in the class that possesses one of the initial n locks and a key ring that is “complete,” *sensu* Bod’ová et al. (2018). That is, they possess the keys to all locks but their own, including the mutant lock. This first class is denoted *L* for “lucky” because its members fortuitously possess the key to the new lock. Each haplotype in this class initially occurs at frequency *d_L_*/*n*, where *d_L_* is an arbitrary initial fraction. Second, for each of these *n_L_* complete haplotypes, there exists one haplotype that is identical except that it lacks the key to the new lock. These could represent ancestral versions of the complete haplotypes that have not yet gained the key to the new lock. This second class is denoted *C* for “chump” because members of this class are simply inferior versions of the corresponding “lucky” haplotypes. Each haplotype in this class occurs at frequency (1 — *d_L_*)/*n*. Third, there are *n_U_* haplotypes that have one of the initial *n* locks and an incomplete key ring that lacks only the key to the mutant lock. Unlike the previous class, there exists no complete version of these haplotypes initially. This third class is denoted *U* for “unlucky” because members of this class cannot unlock the new lock like the “lucky” haplotypes can, but neither must they compete with superior version of themselves like the “chump” haplotypes must. Each haplotype in this class occurs at frequency 1/*n*. Fourth, there is the ancestral haplotype on which the lock mutation arose. It is like any other incomplete haplotype (e.g., a *U* haplotype) except that its key ring is identical to that of the haplotype bearing the lock mutation. This fourth class is denoted *A* for “ancestral” because it is ancestral to the lock mutant. It occurs at frequency (1 – *d_S_*)/*n*, where *d_S_* is the arbitrary initial fraction of ancestral haplotypes that have been replaced with lock-mutant haplotypes. Fifth, there is the haplotype bearing the lock mutation. Unlike all other incomplete haplotype classes, the key that it lacks is the key to its ancestor rather than the key to the new lock. It is denoted *S* for “spiteful” because the lock mutation does not increase the haplotype’s own fitness, but it does decrease the fitness of the “unlucky” and “chump” haplotypes. (Strictly, this is a case of ammensalism rather than spite.) There are thus *n_L_* + *n_U_* + 1 = *n* pre-existing haplotypes (including the mutant’s ancestor) along with the mutant itself for a total of *n* + 1 haplotypes.

**Figure 1:**
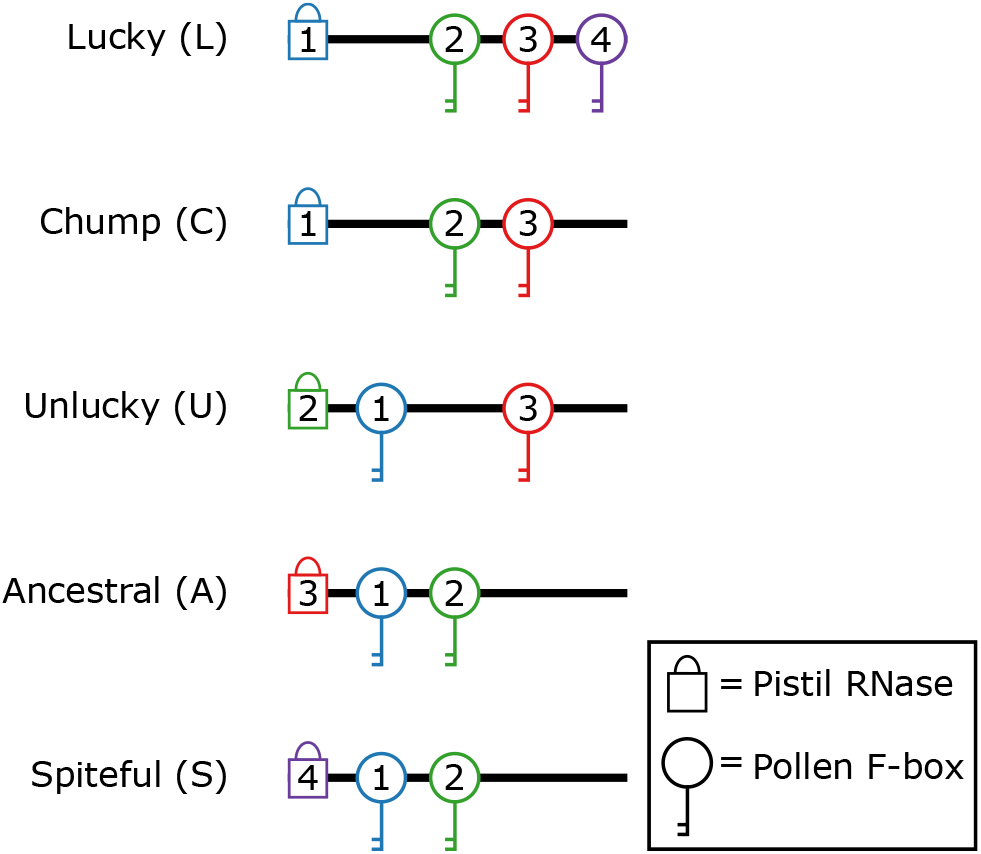
Haplotype classes. In this example, the population begins with three complete haplotypes, and then a fourth lock allele is introduced. The new lock arises from the “ancestral” haplotype, creating a “spiteful” haplotype which is neutral itself but decreases the fitness of the other haplotypes. All haplotypes lack the key to the new lock except for those “lucky” enough to carry it in advance, which have an advantage over the “chump” haplotypes that bear their corresponding locks. Haplotypes that are “unlucky” merely lack the new key but don’t compete against other haplotypes with their same locks.

**Figure 2:**
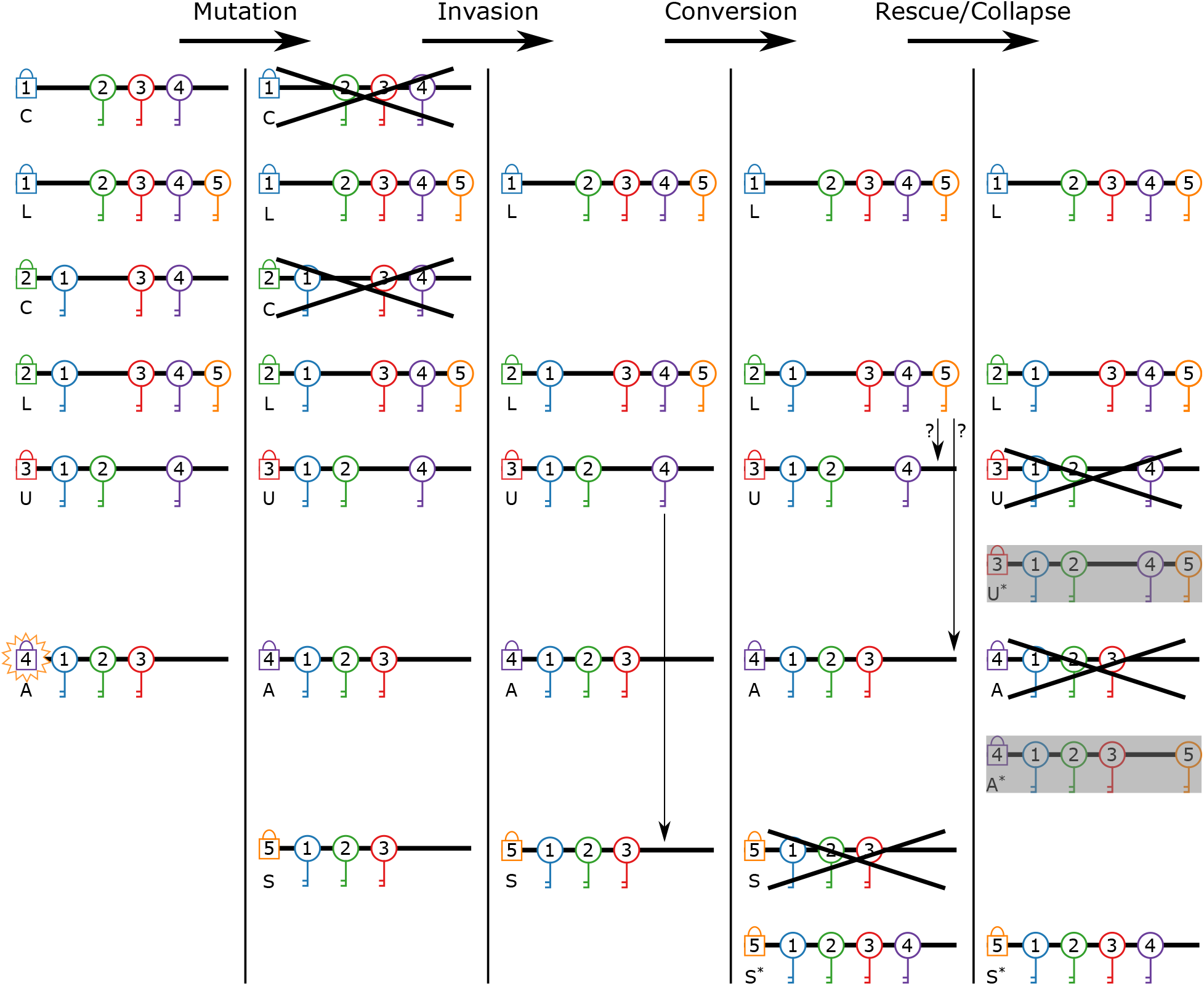
Model steps. These four steps describe a hypothetical pathway to expansion in the number of incompatibility haplotypes. In the “mutation” step, a lock allele mutates to a new specificity (from four to five in this case), one that can be unlocked by an existing, previously neutral key variant. In the “invasion” step, haplotypes with the key to the new lock spread and replace otherwise identical haplotypes lacking the key. In the “conversion” step, haplotypes acquire new keys. This step ends when either the lock mutant acquires the key to its ancestor’s lock, or when its ancestor acquires the key to the new lock. In the “rescue/collapse” step, the remaining haplotypes lacking the key to the new lock are driven to extinction unless they can acquire the missing key in their remaining time. Vertical arrows indicate gene conversion events. Each * represents a haplotype that has acquired a missing key through gene conversion. Question marks indicate gene conversion events that may or may not produce a surviving gene convertant haplotype before the potential recipient is driven extinct. Gray boxes indicate haplotypes resulting from evolutionary rescue events that may or may not occur.

These classes occur at initial frequencies,

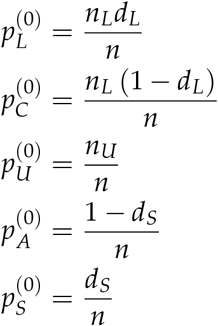

where each *p* represents the frequency of all haplotypes of a given class. That is, the population begins with lock alleles at equal frequencies (as expected at equilibrium before the origin of the new lock) except that a fraction *d_S_* of copies of *A* are replaced by copies of *S*, which induces the distinction between the *L* and *C* classes.

### Invasion of key to novel lock

The population has now passed through the “mutation” step, the first of the four-step pathway (fig. 2). The existence of the new lock fundamentally changes the fitness function in the population. Previously, pollen of each haplotype was equally compatible with all other haplotypes. After the new lock arises, pollen with the new key has an advantage over pollen without it. The fitness gained by a haplotype for being compatible with a particular diploid genotype depends on the frequency of competitors for that genotype. The ability to fertilize a genotype is more advantageous when fewer other haplotypes can also fertilize it. At the instant the lock allele arises, all individuals other than those carrying the new lock allele can be fertilized by the same fraction of the pollen pool,

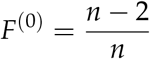

because only two haplotypes (those matching one of the pollen recipient’s pair) are incompatible as sires with a given pollen recipient. In contrast, individuals carrying *S* possess the new lock and can only be fertilized by some fraction of *L* pollen,

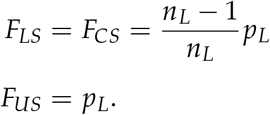

where the subscript indicates a diploid genotype. Genotypes *A A*, *AS* and *SS* do not exist because the *A* and *S* haplotypes are mutually cross-incompatible as well as self-incompatible.

Since all individuals are assumed to be equally fecund as ovule parents and equally viable, all variation in fitness is determined by siring success. The siring success of a given haplotype on a given dam genotype is equal to the frequency of the dam genotype times the frequency of the sire haplotype among all pollen compatible with the dam. We therefore define *D_XY_* of diploid genotype *XY* as the frequency of *XY* divided by the fraction of pollen compatible with *XY*. Comparing marginal fitnesses, *w*,

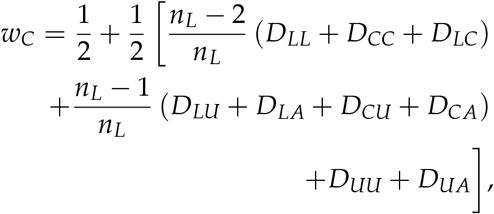

where each *P* represents the frequency of a diploid genotype defined in terms of the two classes of haplotypes it possesses. An *L* haplotype can fertilize everything the corresponding *C* haplotype can, and it can also fertilize individuals carrying the *S* haplotype, so

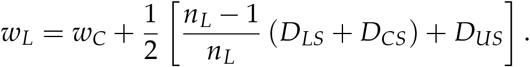

Although *U* haplotypes are compatible with all *L* haplotypes, they are otherwise like *C* haplo-types:

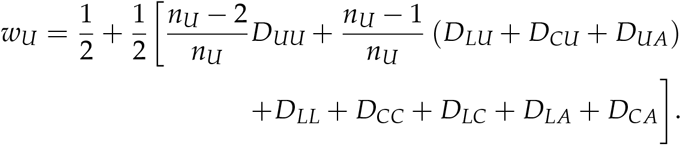

The *A* and *S* haplotypes have equal fitnesses because, although they have different locks, they have identical key rings and thus identical siring prospects:

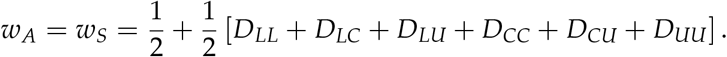

We find that *w_L_ > w_C_* as long as *D_LS_*, *D_CS_*, or *D_US_* > 0 (i.e., as long as *S* exists). That is, as long as *L* haplotypes have a functional key that *C* haplotypes lack, *L* haplotypes have greater fitness. When *S* does not exist, *w_C_* = *w_L_*. Therefore, *w_C_ < w_L_* as long as *S* persists throughout, and all *C* haplotypes are lost at equilibrium. At the initial allele frequencies, the fitnesses *w_A_* and *w_S_* conveniently reduce to

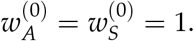

Allele *S* initially invades if 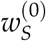 is greater than the initial mean fitness across the haplotypes, 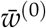, which is true if

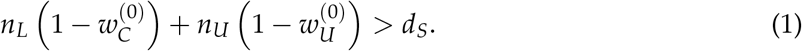

This condition essentially expresses the requirement that there is a sufficient number (*n_L_* or *n_U_*) of sufficiently unfit (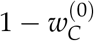 or 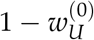) haplotypes such that the lock mutation benefits from their decrease in frequency, and that this quantity must be greater the more copies of the lock mutant there are initially (*d_S_*). Numerical iteration suggests that this condition is not stringent: we considered values of *n_L_, n_U_* ≥ 2 and always found the lock mutation to increase at least slightly.

### Equilibration of haplotype frequencies

The genotype frequency recursion equations (see Appendix) are not linear functions of genotype frequencies, so we cannot derive a vector of equilibrium genotype frequencies by simply solving for an eigenvector of a recursion matrix. Instead, we numerically iterate the genotype frequency recursions for 4000 generations to generate the equilibrium genotype frequencies given *n_U_*, *n_L_, d_L_*, and *d_S_*. We held *d_L_* and *d_S_* both constant at 0.01, and varied *n_U_* and *n_L_* through all combinations of the values 2, 5, 10, and 20 haplotypes. For all these scenarios, the haplotype classes *L, A*, and *S* increased in frequency, while classes *C* and *U* decreased. A representative example with *n_L_ = n_U_* = 2 is plotted in figure 3 for the first 1000 generations. Class *C* approached extinction, and almost all gains went to class *L*. Classes *A* and *S* increased only slightly in frequency.

**Figure 3:**
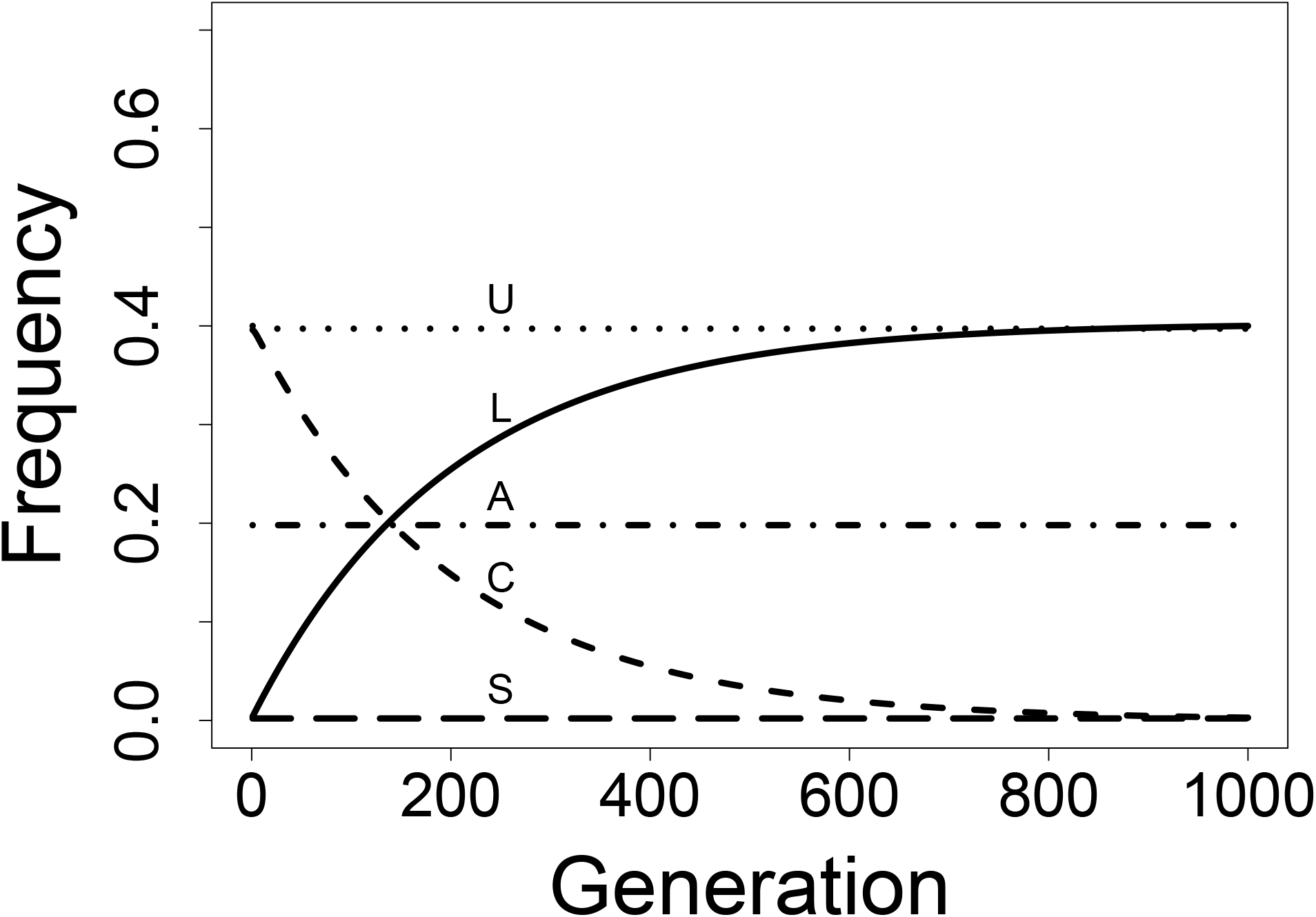
Invasion of a new key. When a new lock (S) arose, any haplotype that fortuitously possessed the key to it (L) invaded and replaced otherwise identical haplotypes lacking the key (C). The lock mutant’s ancestor (A) was unaffected by the lock mutation because it could not fertilize the mutation’s precursor (the ancestor’s own lock) in the first place. The lock mutant (S) had identical siring success as its ancestor, so it was essentially a neutral variant and did not increase in frequency. All other haplotypes (U) decreased slightly in frequency because, with the origin of the new lock, the fraction of individuals they could fertilize decreased slightly. All trajectories were generated by numerical iteration with parameters *d_L_* = *d_S_* = 0.01 and *n_L_* = *n_U_* = 2.

### Acquiring the new key through gene conversion

The population has now passed through the “invasion” step, the second in the four-step pathway (fig. 2). At this point, only a fraction of the population can unlock the new lock. Furthermore, the haplotype with the new lock lacks the key to its ancestor’s lock. But a fully functional haplotype is one that, in pollen, allows fertilization of all plants lacking the same haplotype and, in styles, only prevents fertilization by pollen carrying the same haplotype. To produce such complete haplotypes, all haplotypes except the one possessing the new lock must acquire the key to that lock, and the haplotype with the new lock must acquire the key to its ancestor’s lock. It would be extremely unlikely for each haplotype to acquire the new key by an independent mutation. The new key is also unlikely to be copied to other key rings through recombination because recombinational exchange is rare within the S-locus. However, the low observed allelic diversity at a given F-box paralog locus suggests that these loci have undergone recent gene conversion events (Kubo et al. 2015). Fujii et al. (2016) have shown that if repeated gene conversion events copy the new key to other key rings in series, the products of gene conversion will replace their ancestors without the new key.

We follow Fujii et al. (2016) in modeling the evolution of cross-compatibility through gene conversion. However, as Bod’ová et al. (2018) showed, adding a new key to a haplotype can in some cases give it a universal advantage over others (i.e., an advantage present at any frequency) and potentially drive the others extinct. In particular, if the *S* haplotype acquires the key to its ancestor’s lock, the resulting haplotype is compatible as a sire with every other haplotype, but only a subset of haplotypes are compatible with it. Numerical iteration shows that such a haplotype will rise to high frequencies and eliminate every haplotype that lacks the key to the new lock (fig. 4). These lost haplotypes include all of the U haplotypes along with the lock alleles they carry. Therefore, the overall process results in a collapse in haplotype number at equilibrium rather than expansion.

**Figure 4:**
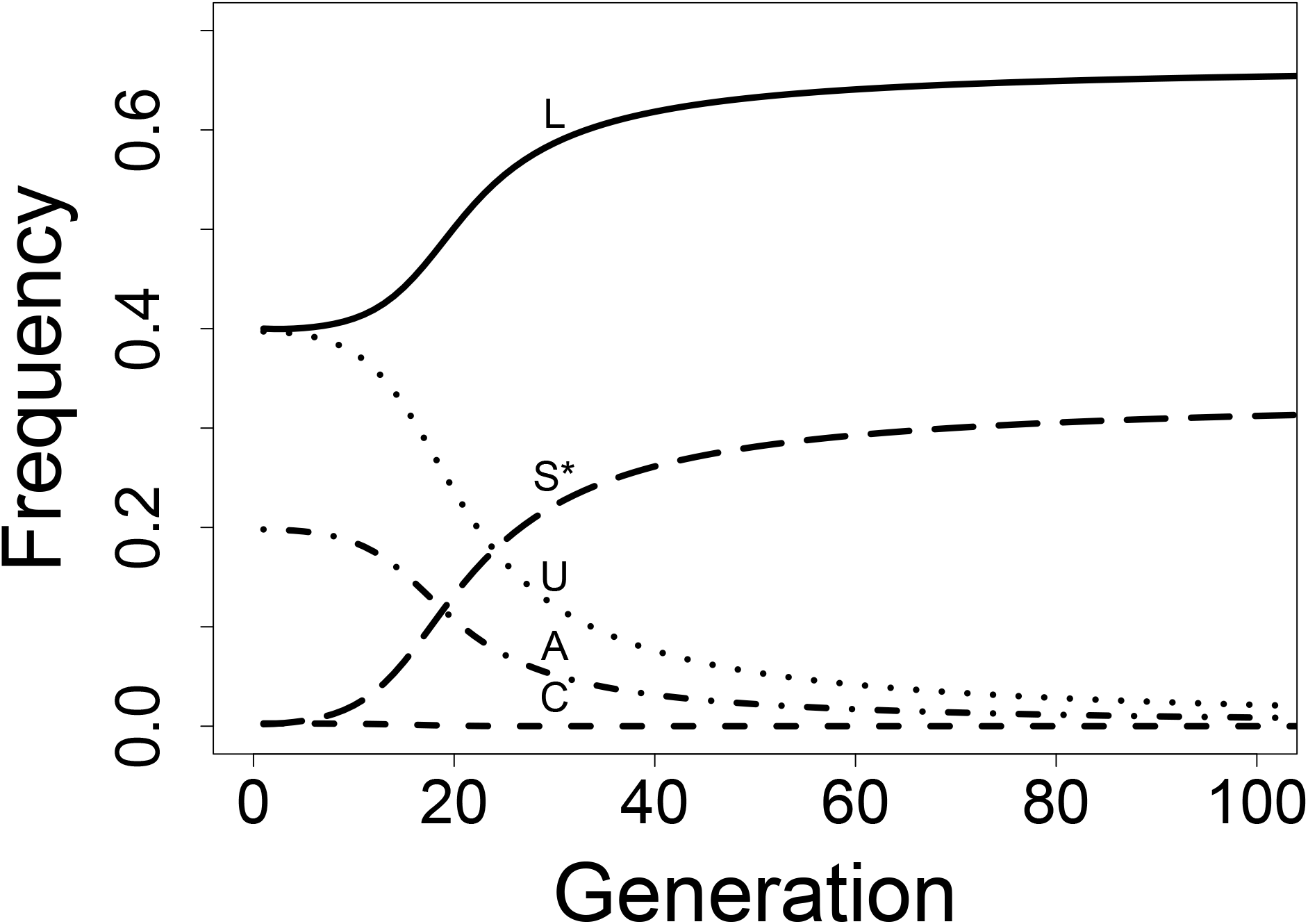
Doomed haplotypes. When the haplotype bearing the lock mutation acquires the key to its ancestor’s lock (becoming S*), it invades. Those haplotypes possessing the key to the new lock (L) also benefit and reach high frequencies, while haplotypes lacking the key (C, U, A) are driven extinct. All trajectories were generated by numerical iteration with parameters *d_L_* = *d_S_* = 0.01 and *n_L_* = *n_U_* = 2.

### Evolutionary rescue of doomed haplotypes

The population has now passed through the “conversion” step, the third in the four-step pathway (fig. 2). Multiple haplotypes are now doomed to extinction at equilibrium. However, if a doomed haplotype acquires the key to the new lock before equilibrium is reached, the resulting haplotype’s fitness increases with the frequency of the new lock. This should protect the resulting gene convertant from being driven from low frequencies to extinction. It may therefore be possible for doomed haplotypes to be rescued by gene conversion. If all haplotypes are rescued before any one of them is lost, then the number of lock alleles at equilibrium is one greater than the initial number, and expansion has occurred. Bod’ová et al. (2018) previously noted the role of evolutionary rescue in this pathway, though they modeled it through recurrent mutation rather than gene conversion. Gene conversion, which Kubo et al. (2015) hypothesized to be responsible for spread of a new key, differs from mutation in several relevant ways. First, whereas a rescuing mutation can occur in any individual carrying a doomed haplotype, functional gene conversion can only occur in individuals carrying both a doomed and a complete haplotype. Gene conversion therefore offers a smaller supply of rescues than does an equal rate of mutation and should provide a lesser probability of rescue. This discrepancy should be minimized when doomed haplotypes, and thus individuals carrying two doomed haplotypes, become rare. There should thus be a form of negative feedback in which the supply of rescuing gene convertants is initially reduced but approaches that of the rescue-by-mutation case as the doomed haplotypes approach extinction. Second, gene conversion does not require recurrent mutations to produce the same key on each doomed haplotype: only a single mutational origin of the rescuing key is necessary. It is conceivable that rescue occurs both through gene conversion and recurrent mutation, but if rescuing mutations are much rarer than rescuing gene conversions, as seems plausible, the supply of rescues may be approximated by the rate of gene conversion alone. Bod’ová et al. (2018) modeled the improbability of recurrent mutation by allowing each key mutation to produce one out of an often large finite set of possible keys, and only those complementary to a currently invading lock could rescue a haplotype. By modeling only gene conversion, we can ignore the larger set of possible keys and instead focus on the rescuing key.

It is also possible that the lock mutant’s ancestral haplotype acquires the key to the new lock before the mutant acquires the key to its ancestor’s lock. The main difference is that, since all other extant haplotypes already possess the key to the ancestor’s lock, only the novel lock must be rescued. Failure to rescue the novel lock will result in a return to the *status quo* rather than a contraction. However, if the novel lock is rescued, the resulting haplotype will begin to exclude the haplotypes lacking its key The process is nearly identical to the case in which the mutant acquires its ancestor’s lock before the ancestor acquires its lock, except that the ancestor is already rescued at the beginning of the process, and the novel lock begins at a lower frequency. We model the invasion of the novel lock, which is necessary for expansion or contraction, but not the rescue of the novel lock, which may or may not occur before its invasion.

The underlying mathematics of this rescue process are similar in form to the evolutionary rescue of a declining population by new mutation. The lock-mutant’s acquisition of its ancestor’s key is a kind of change in the (genetic) environment, the declining frequency of the doomed haplotype is analogous to a declining population, and the acquisition of the new key by the doomed haplotypes is analogous to the generation of new beneficial mutations. We therefore base our model on an existing model of evolutionary rescue by new mutations (Orr and Unckless 2008). There are two components of this model: the supply of beneficial mutations and the probability that any one of them will survive. The probability that the population is rescued is the probability that at least one of these mutations survives. Similarly, the probability that a haplotype is rescued is the probability that at least one copy adds the new key to its key ring and survives. Orr and Unckless (2008) used a model of survival probability in a population of decreasing size (Otto and Whitlock 1997), which is itself a modification of the branching-process model used by Fisher (1923) and Haldane (1927). However, we modify the original model in a different way because, although the declining frequency of the doomed haplotype affects the supply of gene convertants in the same way as a declining population, the effect on survival probability is different. The declining frequency alters the frequency-dependent fitness of the potentially rescuing gene convertants and thus their survival probability. Rather than varying the population size over time, we vary the selection coefficient as a function of genotype frequencies.

In the original model (Orr and Unckless 2008), selection for the rescuing mutation is unconditionally positive. Survival thus implies eventual fixation, and the survival probability equals the fixation probability. In contrast, self-incompatibility involves balancing selection among haplotypes, so survival does not imply fixation. But since the branching-process approximation is actually the probability of surviving early drift, it can be used in either case. Haldane defined *s* as the number of additional offspring produced by the mutant haploid individual above the population average. In our model, *s* can be re-interpreted as the additional number of descendant copies of a gene convertant above the average for all haplotypes. Calculating the relative difference between the marginal fitness of the gene convertant, *w_R_*, and the mean fitness, 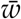, which we define as the selection coefficient *s*,

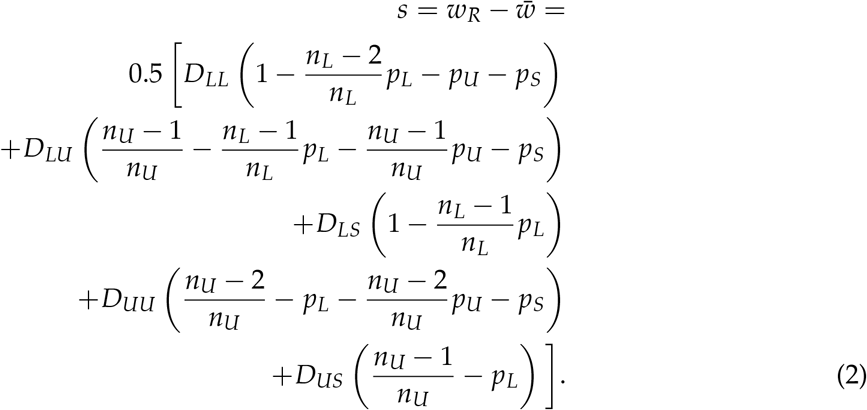

The survival probability for a single gene convertant is thus frequency-dependent, which complicates the calculation of the probability that at least one gene convertant survives.

In the model of Orr and Unckless (2008), the population declines at a predictable rate determined by a standard model of negative population growth. This allowed them to express the number of mutations per generation as a function of time. We cannot do the same with the number of gene conversions per generation because we do not know the genotype frequencies as a function of time. We can, however, numerically iterate recursion equations to get a trajectory of genotype frequencies. We can then retroactively calculate the expected number of gene conversions each generation in this hypothetical history as a product of the per-individual gene conversion rate, the total number of individuals, and the frequency of heterozygotes for the key to the new lock.

Population size and gene conversion rate have no effect on the survival or rescue probability except through their product, so this product is reported as a single compound parameter *R*_conversion_ (conversion-individuals per heterozygote per generation: the exact number of conversions per generation depends on the number of heterozygotes for the key). Note that we use deterministic approximations for the genotype frequency trajectory and the supply of potentially rescuing gene convertants each generation. All stochasticity arises from whether each potentially rescuing convertant actually survives.

The probability of rescue depends on the probability of survival of a new gene convertant. We numerically approximated the survival probability of a new gene convertant using

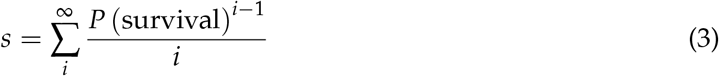

from Haldane (1927). The survival probability of a hypothetical gene convertant was approximated through the following procedure. Each generation, we chose a survival probability that, when substituted into equation (3) (truncated to the first 100 terms of the sum), resulted in a selection coefficient closest to the value calculated through equation (2). Proposed survival probabilities were taken from the sequence from 0 to 1 in increments of 0.01. We checked this approximation against simulations for each generation of a common genotype frequency trajectory with *n_L_* = 2, *n_U_* = 2, and *N* = 10,000. The actual survival probability was determined by simulating the survival of a new gene convertant conditional on arising in a given generation. The convertant was judged to have survived if it survived for at least ten generations. The approximation closely followed the simulated results (fig. S1), so we used the approximation for all future calculations.

Once the genotype frequencies, the expected number of gene convertants, and the survival probability of a new gene convertant were known for every generation, we calculated the probability that none of them survived as the product of the complements of the survival probability. The complement of this probability is the probability that one doomed haplotype is rescued, assuming the expected number of gene convertants under the expected genotype frequency trajectory. We approximated the probability that a given number of haplotypes were rescued as the probability that each of them was rescued independently. The rescue probabilities are not strictly independent because one rescue event alters the trajectory of genotype frequencies. However, they may be approximately independent if the rescue events occur in a relatively short period before genotype frequencies can be greatly altered. Orr and Unckless (2008) found that rescue is most likely to occur early in the process when mutation supply is highest, so it is plausible that rescues also occur in a short period in our model. This procedure resulted in a probability distribution of the number of surviving haplotypes after rescue or collapse. A natural question is, are large collapses guaranteed, or is there a non-trivial probability of expansion?

For all iterations, we assumed that at least two haplotypes had already acquired the key to the new lock. This is the minimum number for the lock-mutation not to confer ovule-sterility because, if only one haplotype could fertilize it, all ovules carrying the lock-mutation would be fertilized by the same pollen haplotype. The maternal haplotype of the resulting offspring would be incompatible with all pollen except the paternal haplotype, which would be incompatible with itself. Therefore, when there is exactly one haplotype of class *L*, all ovules carrying the lock mutation grow up to be ovule-sterile adults. We calculated the probability that all haplotypes survived for all combinations of *n_L_* = 2,5,10,20, *n_U_* = 2,5,10,20, and *R*_conversion_ in the range of 0. 1 – 1 in increments of 0.01 and the range of 1 – 10 in increments of 0.1.

We found that the probability that all haplotypes were rescued (the expansion probability) decreased with the number of doomed haplotypes and increased with the population rate of gene conversion *R*_conversion_ (fig. 5). For *n_L_* = 2, the expansion probability rapidly saturated near 1.0 as *R*_conversion_ increased regardless of *n_U_*. The expansion probability greatly decreased for *n_L_* = 3 relative to *n_L_* = 2 and continued decreasing gradually for larger values of *n_L_*. The expansion probability never reached 0.5 for *n_L_* ≥ 3. The effect of *n_U_*, which was to decrease the expansion probability, was larger for larger *n_L_*.

**Figure 5:**
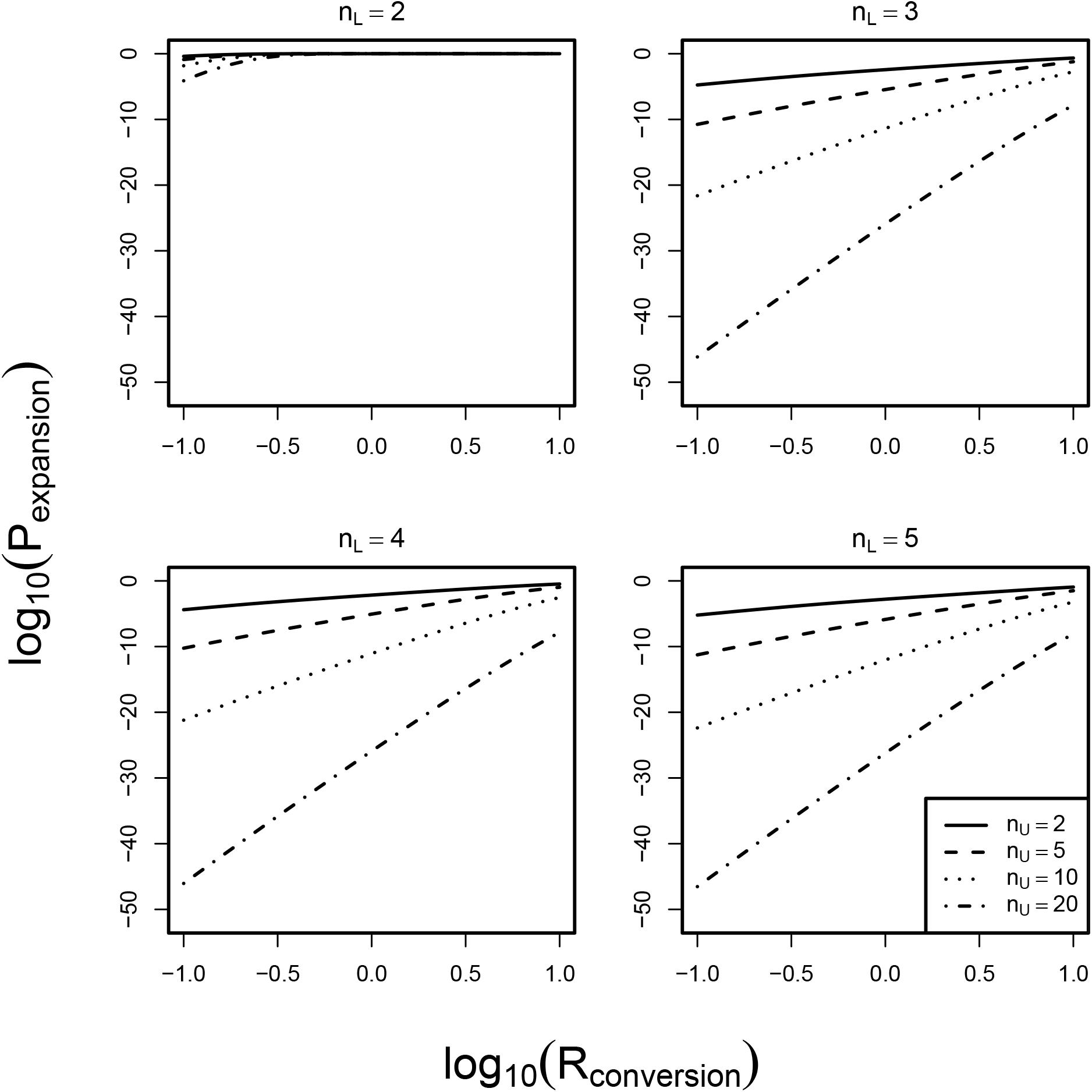
Expansion probability. Each curve represents the probability all doomed haplotypes are rescued for a given number of doomed haplotypes. The probability that all are rescued decreases as the number of doomed haplotypes (*n_U_*) increases, and it also usually decreases as the number of “lucky” haplotypes increases. Rescue is very unlikely unless either the population rate of gene conversion (*R*_conversion_) is very high or *n_L_* is small.

The distribution of haplotype number after rescue/collapse responded similarly to *R*_conversion_ for different values of nL, gradually shifting rightward toward maintenance of the initial number of haplotypes, but never reaching high probabilities of expansion for *R*_conversion_ ≤ 0.1 (fig. 6). For *n_L_* = 2 and *n_L_* = 5, the modal outcome was loss of all or almost all doomed haplotypes.

**Figure 6:**
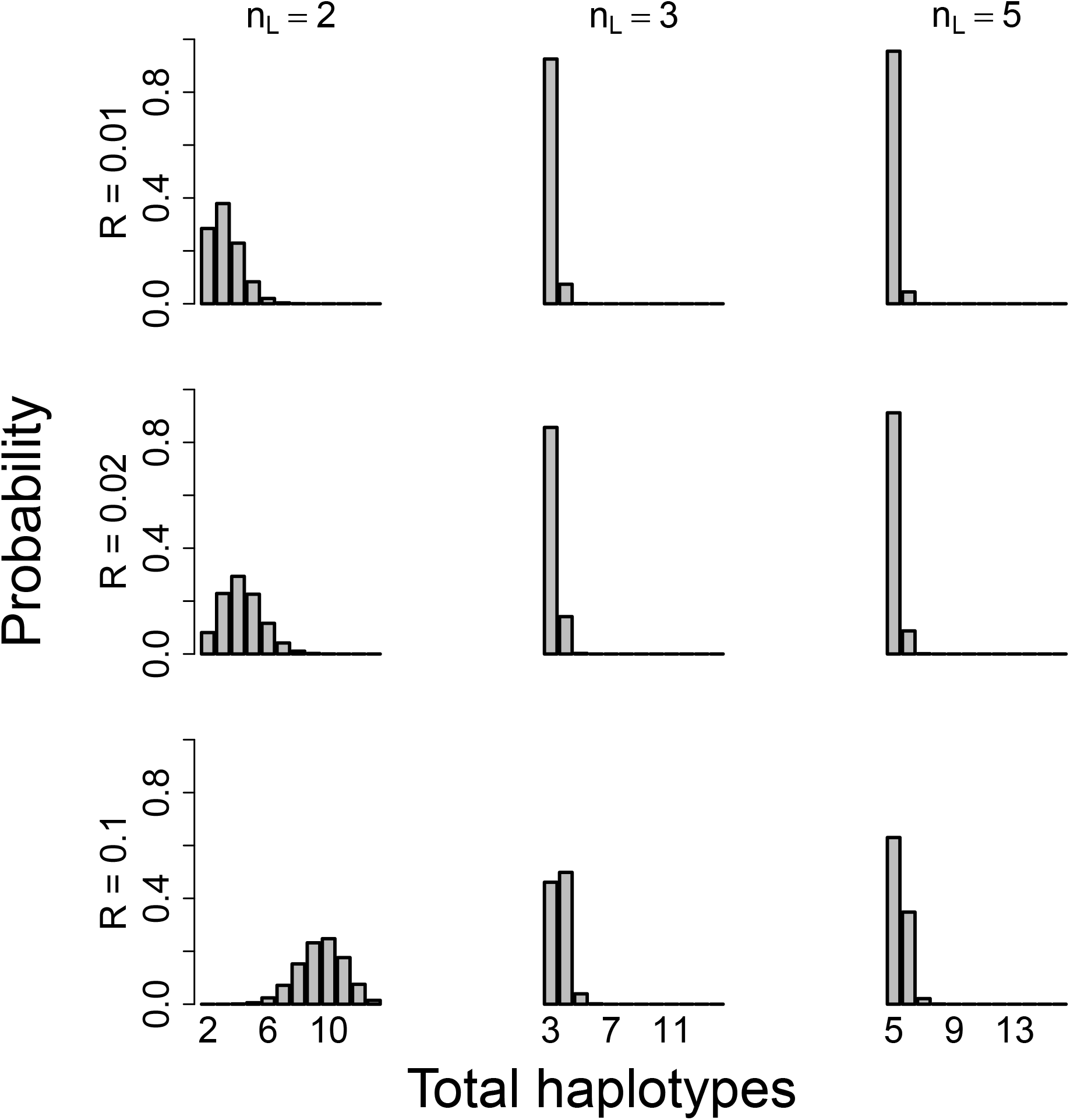
Distribution of final haplotype number. Expansion only occurs when the final number of haplotypes is *n* + 1 (the rightmost bin in each histogram). For *n_L_* > 2 and small population gene conversion rates (*R*_conversion_), the most likely outcomes is either the loss of all doomed haplotypes or the loss of all but one. For all panels, *n_U_* = 10.

### Long-term behavior

The population has now passed through the final “rescue/collapse” step of the four-step pathway (fig. 2). However, natural populations may traverse this pathway repeatedly, and they must have undergone successive rounds of expansion to produce their current haplotypic diversity. The distribution of haplotype number in nature should therefore be governed by the long-term balance between collapse and expansion. We can make crude predictions for the long-term evolution of haplotype number by treating different haplotype numbers as states in a Markov chain. The transition process works as follows. In the initial state, there are *n* = *n_L_* + *n_U_* + 1 unique lock alleles, including the one carried by the single haplotype of class A. After the new lock invades, there are *n* + 1 locks, now also including that of the single haplotype of class *S*. After the rescue/collapse phase, the new state is the number of surviving lock alleles, ranging from *n*’ = *n* – *n_U_* = *n_L_* + 1 (no locks are rescued; only those sharing a haplotype with a complete key ring survive) to *n*’ = *n* + 1 = *n_L_* + *n_U_* + 2 (all locks are rescued). Assuming that haplotype number changes only through the foregoing pathway, the stationary distribution of this Markov chain represents the probability distribution for haplotype number at the expansion-collapse equilibrium. Other unmodeled processes, such as bottlenecks or alternative diversification pathways, may decrease or increase the number observed in nature relative to the values predicted by the Markov chain.

At the end of one transition, all haplotypes possess the key to the lock that just invaded. However, this does not imply anything about whether they possess the key to the next lock to invade. The haplotypes that were lucky with respect to one lock may be unlucky with respect to the next. The outcome of the rescue process therefore tells us *n*’ but not 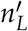 or 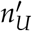 individually. We assume that *n_L_* remains a constant parameter between states, and that all changes in haplotype number are represented by changes in *n_U_*.

Our rescue probability calculations showed that expansion was likely even from large initial haplotype numbers when *n_L_* = 2 and *R*_conversion_ ≥ 0.1 (fig. 5). Such large expansion probabilities are necessary to produce the many haplotypes observed in nature. To cover a range of expansion probabilities, we examined *R*_conversion_ = 0.1–0.9 in intervals of 0.1, as well as *R*_conversion_ = 0.01 and 1. The low value of *n_L_* = 2 is biologically reasonable because an initially neutral key would likely be present on few key rings. (A value of *n_L_* = 1 may be even more common, but it would impose a severe penalty on the new lock: *L_S_* would become the only S-bearing genotype and would be ovule-sterile.)

The time steps in this Markov process are somewhat abstract: they represent the waiting times between the invasion of one lock and the next. In reality, they would be variable in length. However, neither rescue nor collapse can occur during these periods. The length of each waiting period is therefore irrelevant to the diversification process we modeled, though it may affect the opportunity for bottlenecks or alternative diversification pathways. Since we were unable to map the waiting periods onto chronological time, we ignored the dynamics of the process and focused on the ultimate stationary distribution. Preliminary trials showed that the final distribution was unaffected by the initial distribution, and we set the initial distribution such that all probability was concentrated at *n_U_* = 1.

Our numerical iterations required a transition matrix of finite size. We therefore enforced an upper boundary of 40 incomplete haplotypes, near the upper range observed in nature (Lawrence 2000). With the fixed value of *n_L_* = 2, this resulted in up to 42 total haplotypes. Any non-zero probability of expanding beyond this maximum was instead treated as additional probability of remaining at the maximum. We also enforced a lower boundary at the biological minimum of *n* = 3: below this value, all plants reject all pollen and the population is completely sterile. Any non-zero probability of collapsing to a state in which *n* < 3 was instead treated as additional probability of transitioning to *n* = 3.

We found that the stationary distribution shifted toward greater numbers of haplotypes as *R*_conversion_ increased (fig. 7). At *R*_conversion_ = 0.01, almost all probability was concentrated at the biological minimum of three haplotypes, while at *R*_conversion_ = 1, almost all probability was concentrated at 40 haplotypes, the maximum we allowed. In the range from *R*_conversion_ = 0.1–0.3, the stable distribution was centered around intermediate values in the range 3–40.

**Figure 7:**
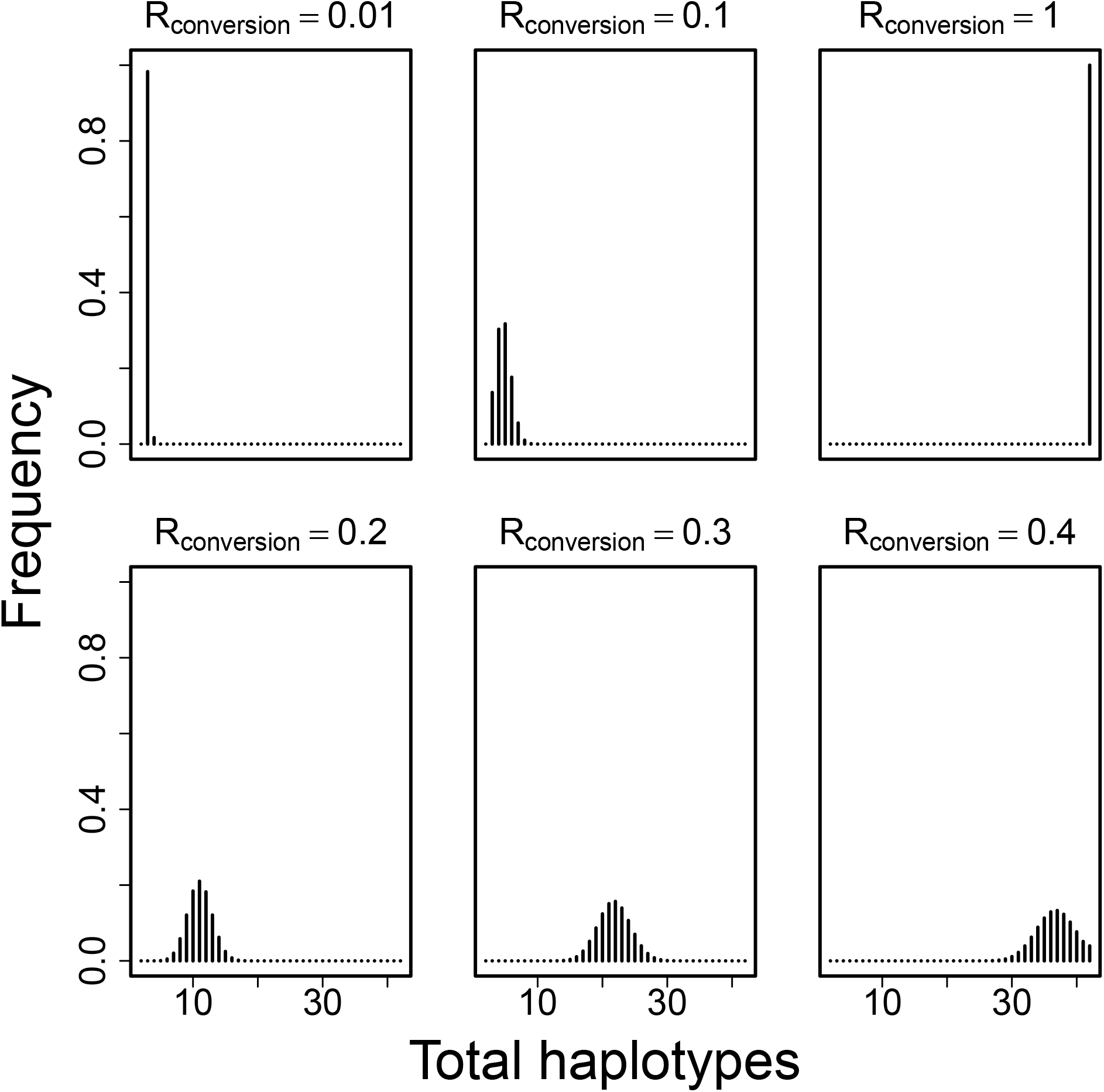
Stable distribution of haplotype number. The stable number of haplotypes increased as *R*_conversion_, the supply of gene convertants, increased. The haplotype number came to exceed the biological minimum near *R*_conversion_ = 0.1, and produced 20–40 haplotypes around *R*_conversion_ = 0.3 or 0.4. For all panels, the Markov chain was run for 1000 steps (expansions or collapses), and *n_L_* = 2. All probability of producing more than 40 incomplete haplotypes (i.e., 42 total haplotypes) is binned into the 42nd category.

## Discussion

We used an evolutionary rescue model to estimate the relative probabilities of expansion and contraction in S-haplotype number, calculate the distribution of contraction magnitudes, and predict the long-term evolution of haplotype number. We find that expansion from low haplotype number is possible if gene conversion is frequent or the population is large, but that the possibility of contraction is ubiquitous. These collapses can be large, easily resulting in the loss of the majority of S-haplotypes. A unique prediction of this model is that recently bottlenecked populations should have a “debt” such that, once the appropriate gene conversion event occurs, they will experience a sudden reduction in haplotype number in addition to the reduction directly caused by the bottleneck. The long-term maintenance of haplotype number is therefore more precarious than previously appreciated.

This vulnerability is a necessary consequence two very reasonable premises along with what is known empirically about the control of self-incompatibility. First, we assumed that gene conversion can copy any F-box specificity from one haplotype to another. This is consistent with the low observed polymorphism among alleles of a given F-box paralog despite their long coexistence (Kubo et al. 2015). Second, we assumed that the supply of compatible pollen did not limit seed set. Pollen limitation certainly occurs in nature: for self-incompatible plants surveyed in the literature, Larson and Barrett (2000) found on average a 59% increase in fruit set for pollen-supplemented flowers relative to open-pollinated flowers, significantly greater than the 31% average for self-compatible species. However, even their set of 66 self-incompatible species contained over 10 species with 0-20% pollen limitation. Self-incompatible populations with little pollen limitation may therefore be moderately frequent, even if they are not the norm.

Despite the possibility of collapse, we found that stable distributions of haplotype number within the range of 20–40 were possible in the long term. These numbers are greater than the upper limit of 14 haplotypes found in the model of Bod’ová et al. (2018), and they are within the range found in nature. This discrepancy is most likely due to the large supplies of rescuing gene convertants we modeled, and it shows that arbitrarily many haplotypes can be maintained if the population is large or gene conversion is rapid. Population size and S-haplotype number have rarely been estimated for the same populations. However, complete sampling of three populations of *Pyrus pyraster,* a rare woody perennial, revealed 9–25 S-haplotypes per population despite small (N = 8–88) population sizes (Hoebee et al. 2011). Our model fails to predict the maintenance of so many haplotypes in such small populations at equilibrium, but it is possible that these populations are not at equilibrium. If the waiting times between successive RNase invasions are long, then it is possible that collapses in haplotype number lag far behind reductions in population size. In this case, we should predict that S-haplotype number will drop precipitously in *P. pyraster* in the (possibly distant) future.

The rate at which gene conversion copies a functional F-box from one haplotype to another is unknown. Per-nucleotide rates of gene conversion have been estimated to be 400 times the rates of crossovers in *Drosophila* (Gay et al. 2007). Assuming a crossover rate of 10^−8^ per meiosis per nucleotide, this yields a gene conversion rate on the order of 10^−6^. A functional gene conversion rate of 10^−6^ would produce few haplotypes (≈ 3) for populations smaller than 100,000, ≈ 10 – 40 for populations on the order of 100,000, and larger numbers for larger populations. Such populations are large but not unrealistic. Note, however, that the parameter in our model is not merely the per-nucleotide rate, but the rate at which a functional F-box is gained. This rate excludes gene conversion events that eliminate rather than transfer a specificity as well as events that fail to affect specificity at all.

The amount of nucleotide divergence between functionally distinct F-box paralogs could serve as an upper bound on how many changes are required to convert one specificity to another. However, this quantity would also include neutral substitutions unrelated to the difference in specificity. A direct but technically difficult means of estimating the rate of functional gene conversion would be to genotype individual pollen grains of a self-incompatible individual using long-read sequencing and to detect haplotypes with the F-box paralog corresponding to their own RNase. Alternatively, a self-incompatible individual could be self-pollinated. Although most pollen would be rejected, some fraction would have become self-compatible through gene conversion. Such spontaneous gene conversion could be distinguished from mere leaky self-incompatibility either through long-read sequencing of the offspring or by further crosses. Gene-convertant offspring should be partially self-compatible because half their pollen possesses the F-box genes complementary to both parental RNases. Gene conversion could be distinguished from self-compatibility mutations in the pistil locus or modifier loci because gene convertants should still fully reject pollen from plants with the same RNase-locus genotype: only their pollen phenotype should change. Although both new mutations and gene conversion could add a pollen specificity and induce self-compatibility, these processes need not be distinguished because their effects are identical from the perspective of haplotype diversification.

Other than large populations are rapid gene conversion, another possible explanation of high S-haplotype diversity is that gene flow buoys species-wide diversity despite local contractions. Uyenoyama et al. (2001) found that, similar to our results, S-allele number was unlikely to expand within a single population because of the threat of collapse into self-compatibility. They proposed that expansion may instead occur through local turnover in S-alleles followed by introduction of the novel alleles into the broader metapopulation. However, this model assumed that incompatibility operates through self-recognition, in which introgressed S-alleles gain an advantage because they are compatible with resident plants that do not recognize them as self. It is thus not obvious whether the conclusion translates to nonself-recognition. Castric et al. (2008) found less divergence between putative shared ancestral variants of S-locus genes in *Arabidopsis lyrata* and *A. halleri* than between non-S orthologs, consistent with elevated introgression at the S-locus in this self-recognition system. Under collaborative nonself-recognition, however, each novel RNase requires a corresponding F-box. A novel introgressed RNase allele will not necessarily be detoxified by local F-box proteins, and it may thus inflict ovule-sterility. If there is insufficient compatible pollen to support the migrant haplotype, self-incompatibility may act as a barrier to introgression not just at the S-locus, but genome-wide. This barrier could be overcome, but it would require either that the key corresponding to the foreign lock already exists in the population (e.g., as a dual-function key or as a segregating neutral variant), or that there is sufficient migration to supply the corresponding foreign keys. Reintroduction of recently lost RNases, unlike introduction of truly novel RNases, should function identically under self- and nonself-recognition. Populations that have undergone a recent contraction meet this condition because they have lost RNases but not necessarily the corresponding F-box genes. As long as the lost RNases were reintroduced from another population before the F-box genes became pseudo-genized, the population could recover all lost S-haplotypes. A metapopulation can therefore still provide a “safety net” that prevents permanent contraction even if truly novel RNases cannot easily spread.

Highly differentiated sets of RNases may even act to generate reproductive isolation. If nonself-recognition poses greater barriers to introgression than self-recognition, this tendency might partly explain species selection for self-incompatibility: clades with nonself-recognition may more rapidly undergo reproductive isolation and thus speciation. Landis et al. (2018) found either a positive effect or no effect of autogamy on net diversification in Polemoniaceae, in contrast to the negative effect of self-incompatibility on net diversification found by Goldberg et al. (2010) in Solanaceae. Although Goldberg et al. (2010) attributed the greater net diversification rate in self-incompatible species to a lesser extinction rate compared to self-compatible species rather than a greater speciation rate, it is difficult to estimate these quantities separately. If the increased diversification rate in the self-incompatible species of Solanaceae is due to isolating effects of nonself-recognition rather than from self-incompatibility *per se*, we should expect a reduced effect of self-incompatibility on diversification in families with other systems like Brassicaceae and Asteraceae.

We found that the expansion probability greatly decreased as the number of complete haplotypes increased. Greater numbers of complete haplotypes reduce the expansion probability in at least two ways. First, by increasing the number and thus the frequency of fit haplotypes, they increase the mean siring success. When compatibility with the new RNase is more common, the competitive advantage of this compatibility is reduced. Second, with more complete haplotypes, the equilibrium frequency of the novel RNase is lesser. These two effects reduce the advantage of compatibility with the new RNase and thus the survival probability of the rescuing gene convertant. Expansion was most likely when exactly two haplotypes were initially complete. This condition may be more biologically plausible than it first appears. Most F-box mutations should exist on only one haplotypic background because all new mutations fall into this class. However, these alleles should contribute neither to expansion nor contraction: they cannot support a corresponding RNase mutation in the first place unless they exist on at least two backgrounds. If F-box mutations on two backgrounds are the second-most-common, then they are the most common within the subset that can actually contribute to contraction or expansion.

An essential parameter of this model, the number of haplotypes pre-adapted to the novel RNase specificity, has not been systematically estimated. The existence of multiple pre-adapted haplotypes is a prerequisite for both expansion and collapse. The presence and number of pre-adapted haplotypes in a population could be determined by experimentally applying naïve pollen to plants carrying a novel, functional RNase allele. Successful fertilization would indicate pre-adaptation to the RNase. An alternative explanation for fertilization, the loss of RNase function in the dam, could be ruled out with the appropriate control. F-box knockouts should be compatible with RNase loss-of-function mutants but not with novel RNase alleles. Novel RNase alleles could be generated by mutagenesis, or they could be introduced from distantly related populations transgenically or through introgression.

Besides bounding the parameters of the model, empirical research could also test for evidence of recent historical collapses. Consider a pair of allopatric populations that initially shared all S-haplotypes. However, a novel RNase has recently invaded Population A and eliminated several pre-existing haplotypes, resulting in a contraction in haplotype number. This RNase never arose in Population B, which retains the original complement. Every remaining haplotype in Population A other than the novel haplotype itself should carry the F-box complementary to the novel RNase. In Population B, several haplotypes should either be polymorphic for the novel RNase’s complementary F-box or should lack it entirely. This situation could be tested by reciprocally crossing individuals from the two populations. A substantial proportion of B pollen should be rejected: a proportion in excess of that predicted by the number and frequency of shared S-haplotypes. A potentially dramatic test would be to introgress the novel RNase from Population A to Population B. In the short run, the siring success of all haplotypes not pre-adapted to the new RNase would decrease. In the long run, the frequency of these unlucky haplotypes would decrease to extinction. That is, we predict that S-haplotype collapse should be contagious between closely related populations, even if both populations are self-incompatible.

SI has developed into a rich study system through the continued interaction between theory and empirical research. Experimental demonstration of the genetic control of SI (East and Mangelsdorf 1925) and field research showing the number of S-alleles in natural populations (Emerson 1938,1939) inspired theoretical explanations of the balancing selection capable of maintaining this diversity (Wright 1939; Charlesworth and Charlesworth 1979). The theoretical potential for balancing selection indicated the S-locus as a candidate for long-term polymorphism, and S-allele phylogenies confirmed this possibility (Igic and Kohn 2001; Steinbachs and Holsinger 2002). The discovery of the fine-scale genetic basis of nonself-recognition (Kubo et al. 2010, 2015) necessitated new theoretical explanation of the expansion process, and this theory now points to the unanticipated possibility of S-allele collapse through runaway gene convertants.

## Appendix: genotype frequency recursions

The following genotype frequency recursions were used to determine the equilibrium genotype frequencies after the invasion of the key to the new lock (fig. 3). They were also used to produce the trajectories leading to extinction of doomed haplotypes (fig. 4), which were in turn used to calculate rescue probabilities. The genotype frequency recursions are

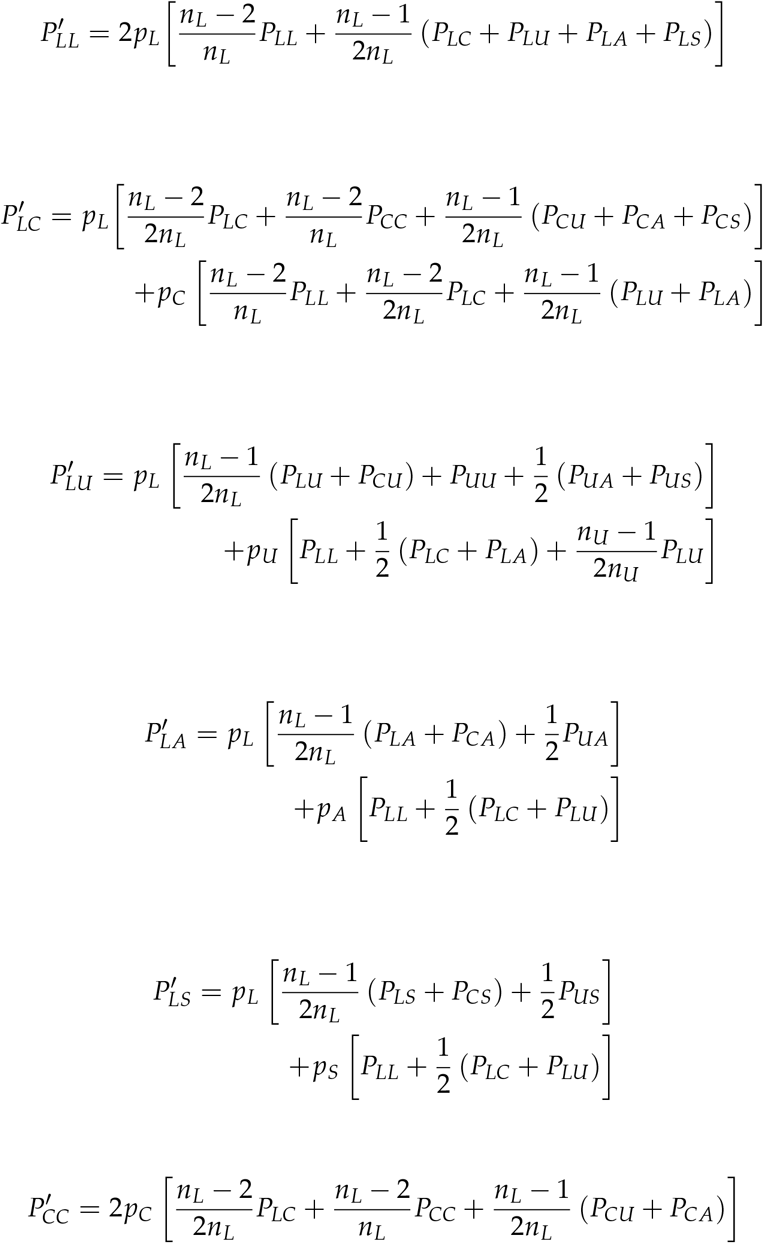

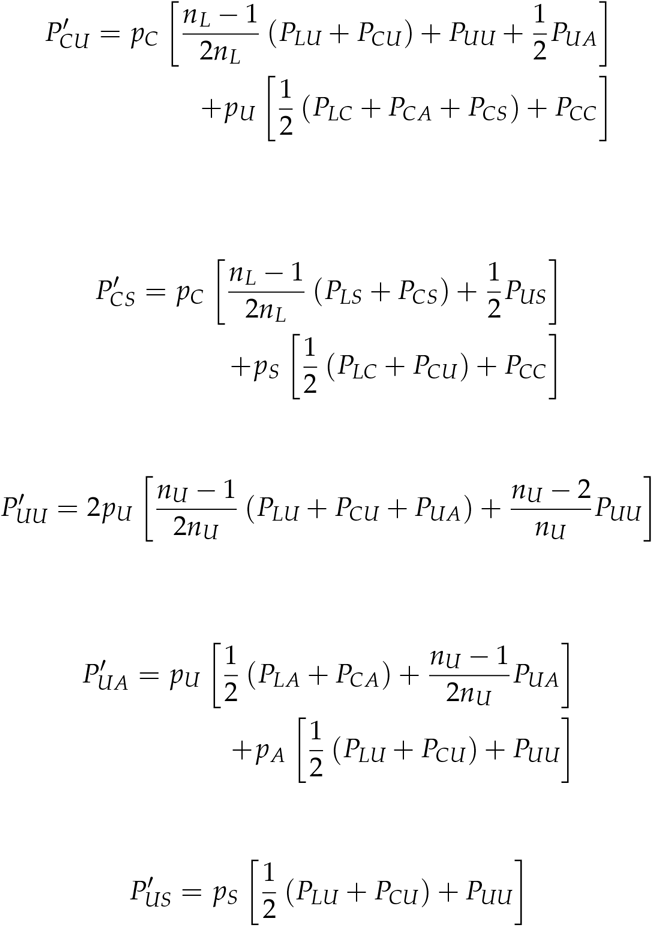

---

**Figure S1:**
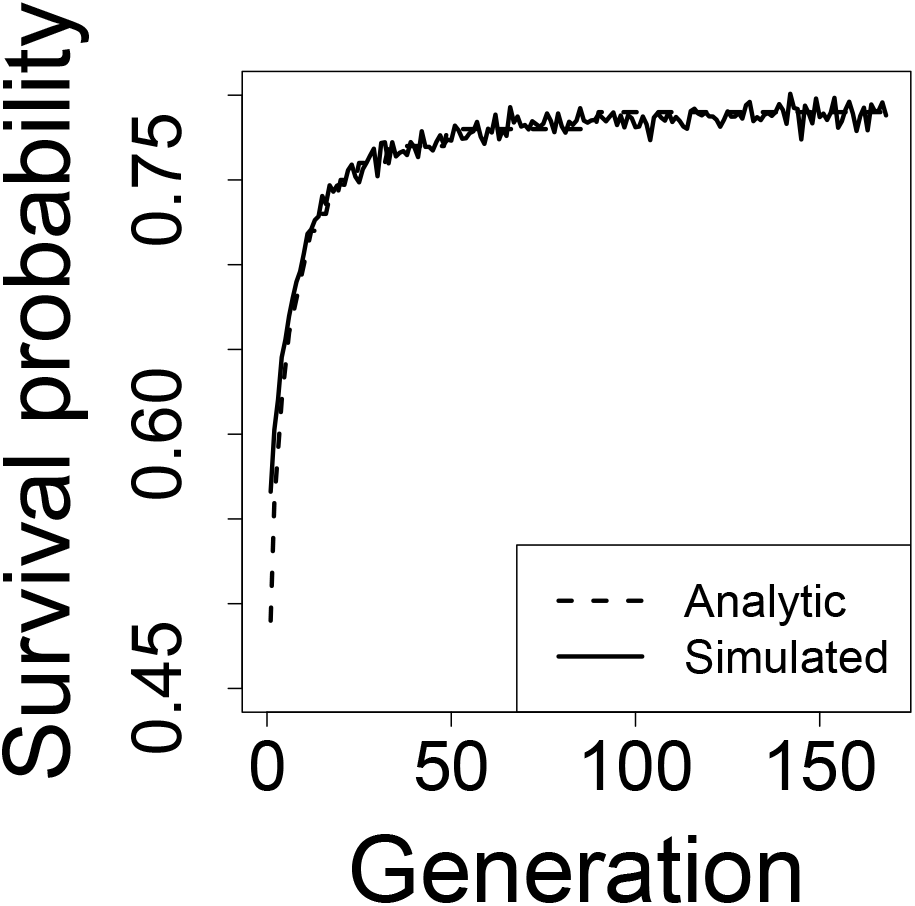
Survival probability of a new gene convertant. The history of a population with *n_L_* = 2, *n_U_* = 2, *N* = 10,000 during the rescue phase was first determined by iterating the recursion equations. Survival probability was determined each generation using Haldane’s series expansion (analytic) or by simulating survival of a new gene convertant for 10 generations over 10,000 replicate simulations (simulated).

## Literature Cited

Bod’ová, K., T. Priklopil, D. L. Field, N. H. Barton, and M. Pickup. 2018. Evolutionary pathways for the generation of new self-incompatibility haplotypes in a non-self recognition system. Genetics 209:861–883.

Castric, V., J. Bechsgaard, M. H. Schierup, and X. Vekemans. 2008. Repeated adaptive introgression at a gene under multiallelic balancing selection. PLoS Genetics 4.

Charlesworth, D., and B. Charlesworth. 1979. The evolution and breakdown of S-allele systems. Heredity 43:41–55.

East, E. M., and A. J. Mangelsdorf. 1925. A new interpretation of the hereditary behavior of self-sterile plants. Proceedings of the National Academy of Sciences of the United States of America 11:166–171.

Emerson, S. 1938. The genetics of self-incompatibility in *Oenothera organensis*. Genetics 23:190.

Emerson, S. 1939. A preliminary survey of the *Oenothera organensis* population. Genetics 24:524.

Entani, T., M. Iwano, H. Shiba, F.-S. Che, A. Isogai, and S. Takayama. 2003. Comparative analysis of the self-incompatibility (S-) locus region of *Prunus mume*: identification of a pollen-expressed F-box gene with allelic diversity. Genes to Cells 8:203–213.

Fisher, R. A. 1923. XXI.—On the dominance ratio. Proceedings of the Royal Society of Edinburgh 42:321–341.

Foote, H., J. P. Ride, V. E. Franklin-Tong, E. A. Walker, M. J. Lawrence, and F. Franklin. 1994. Cloning and expression of a distinctive class of self-incompatibility (S) gene from *Papaver rhoeas* L. Proceedings of the National Academy of Sciences 91:2265–2269.

Fujii, S., K.-i. Kubo, and S. Takayama. 2016. Non-self-and self-recognition models in plant self-incompatibility. Nature Plants 2.

Gay, J. C., S. Myers, and G. McVean. 2007. Estimating meiotic gene conversion rates from population genetic data. Genetics 177:881–894.

Gervais, C. E., V. Castric, A. Ressayre, and S. Billiard. 2011. Origin and diversification dynamics of self-incompatibility haplotypes. Genetics 188:625–636.

Goldberg, E. E., J. R. Kohn, R. Lande, K. A. Robertson, S. A. Smith, and B. Igić. 2010. Species selection maintains self-incompatibility. Science 330:493–495.

Haldane, J. B. S. 1927. A mathematical theory of natural and artificial selection, part V: selection and mutation. Pages 838–844 *in* Mathematical Proceedings of the Cambridge Philosophical Society. Vol. 23. Cambridge University Press.

Hiscock, S. J., S. M. McInnis, D. A. Tabah, C. A. Henderson, and A. C. Brennan. 2003. Sporophytic self-incompatibility in *Senecio squalidus* L. (Asteraceae)—the search for S. Journal of Experimental Botany 54:169–174.

Hoebee, S., S. Angelone, D. Csencsics, K. Määttänen, and R. Holderegger. 2011. Diversity of S-alleles and mate availability in 3 populations of self-incompatible wild pear *(Pyrus pyraster)*. Journal of Heredity 103:260–267.

Igić, B., and J. R. Kohn. 2001. Evolutionary relationships among self-incompatibility RNases. Proceedings of the National Academy of Sciences 98:13167–13171.

Igić, B., R. Lande, and J. R. Kohn. 2008. Loss of self-incompatibility and its evolutionary consequences. International Journal of Plant Sciences 169:93–104.

Kubo, K.-i., T. Entani, A. Takara, N. Wang, A. M. Fields, Z. Hua, M. Toyoda, S.-i. Kawashima, T. Ando, A. Isogai, et al. 2010. Collaborative non-self recognition system in S-RNase–based self-incompatibility. Science 330:796–799.

Kubo, K.-i., T. Paape, M. Hatakeyama, T. Entani, A. Takara, K. Kajihara, M. Tsukahara, R. Shimizu-Inatsugi, K. K. Shimizu, and S. Takayama. 2015. Gene duplication and genetic exchange drive the evolution of S-RNase-based self-incompatibility in *Petunia*. Nature Plants 1.

Lai, Z., W. Ma, B. Han, L. Liang, Y. Zhang, G. Hong, and Y. Xue. 2002. An F-box gene linked to the self-incompatibility (S) locus of *Antirrhinum* is expressed specifically in pollen and tapetum. Plant Molecular Biology 50:29–41.

Landis, J. B., C. D. Bell, M. Hernandez, R. Zenil-Ferguson, E. W. McCarthy, D. E. Soltis, and P. S. Soltis. 2018. Evolution of floral traits and impact of reproductive mode on diversification in the phlox family (Polemoniaceae). Molecular Phylogenetics and Evolution 127:878–890.

Larson, B. M., and S. C. Barrett. 2000. A comparative analysis of pollen limitation in flowering plants. Biological Journal of the Linnean Society 69:503–520.

Lawrence, M. 2000. Population genetics of the homomorphic self-incompatibility polymorphisms in flowering plants. Annals of Botany 85:221–226.

Li, X., N. Paech, J. Nield, D. Hayman, and P. Langridge. 1997. Self-incompatibility in the grasses: evolutionary relationship of the S gene from *Phalaris coerulescens* to homologous sequences in other grasses. Plant Molecular Biology 34:223–232.

McClure, B. A., V. Haring, P. R. Ebert, M. A. Anderson, R. J. Simpson, F. Sakiyama, and A. E. Clarke. 1989. Style self-incompatibility gene products of *Nicotiana alata* are ribonucleases. Nature 342:955–957.

Orr, H. A., and R. L. Unckless. 2008. Population extinction and the genetics of adaptation. The American Naturalist 172:160–169.

Otto, S. P., and M. C. Whitlock. 1997. The probability of fixation in populations of changing size. Genetics 146:723–733.

Paape, T., B. Igic, S. D. Smith, R. Olmstead, L. Bohs, and J. R. Kohn. 2008. A 15-myr-old genetic bottleneck. Molecular Biology and Evolution 25:655–663.

Sakai, S. 2016. How have self-incompatibility haplotypes diversified? Generation of new haplotypes during the evolution of self-incompatibility from self-compatibility. The American Naturalist 188:163–174.

Sijacic, P., X. Wang, A. L. Skirpan, Y. Wang, P. E. Dowd, A. G. McCubbin, S. Huang, and T.-h. Kao. 2004. Identification of the pollen determinant of S-RNase-mediated self-incompatibility. Nature 429:302–305.

Stein, J. C., B. Howlett, D. C. Boyes, M. E. Nasrallah, and J. B. Nasrallah. 1991. Molecular cloning of a putative receptor protein kinase gene encoded at the self-incompatibility locus of *Brassica oleracea*. Proceedings of the National Academy of Sciences 88:8816–8820.

Steinbachs, J., and K. Holsinger. 2002. S-RNase–mediated gametophytic self-incompatibility is ancestral in eudicots. Molecular Biology and Evolution 19:825–829.

Tank, D. C., J. M. Eastman, M. W. Pennell, P. S. Soltis, D. E. Soltis, C. E. Hinchliff, J. W. Brown, E. B. Sessa, and L. J. Harmon. 2015. Nested radiations and the pulse of angiosperm diversification: increased diversification rates often follow whole genome duplications. New Phytologist 207:454–467.

Uyenoyama, M. K., Y. Zhang, and E. Newbigin. 2001. On the origin of self-incompatibility haplotypes: transition through self-compatible intermediates. Genetics 157:1805–1817.

Wright, S. 1939. The distribution of self-sterility alleles in populations. Genetics 24:538–552.

